# Cell-surface PD-L1 expression identifies a sub-population of distal epithelial cells enriched in idiopathic pulmonary fibrosis

**DOI:** 10.1101/2022.03.09.483616

**Authors:** Negah Ahmadvand, Gianni Carraro, Matthew R. Jones, Irina Shalashova, Jochen Wilhelm, Nelli Baal, Farhad Kosravi, Chengshui Chen, JinSan Zhang, Clemens Ruppert, Andreas Guenther, Roxana Wasnick, Saverio Bellusci

## Abstract

Idiopathic lung fibrosis (IPF) is a fatal lung disease characterized by chronic epithelial injury and exhausted repair capacity of the alveolar compartment, associated with the expansion of cells with intermediate alveolar epithelial cell (AT2) characteristics. Using *Sftpc^CreERT2/+^: tdTomato^flox/flox^* mice, we previously identified a lung population of quiescent injury activated alveolar epithelial progenitors (IAAPs) marked by low expression of the AT2 lineage trace marker tdTomato (Tom^low^), and characterized by high levels of Pd-l1 (Cd274) expression. This led us to hypothesize that a population with similar properties exists in the human lung. To that end, we used flow cytometry to characterize the CD274 cell surface expression in lung epithelial cells isolated from donor and end-stage IPF lungs. The identity and functional behavior of these cells were further characterized by qPCR analysis, *in vitro* organoid formation, and *ex vivo* precision-cut lung slices (PCLS). Our analysis led to the identification of a population of CD274^pos^ cells expressing intermediate levels of *SFTPC*, which was expanded in IPF lungs. While donor CD274^pos^ cells initiated clone formation, they did not expand significantly in 3D organoids in AT2-supportive conditions. However, an increased number of CD274^pos^ cells was found in cultured PCLS. In conclusion, we demonstrate that, similar to the IAAPs in the mouse lung, a population of CD274 expressing cells exists in the normal human lung and this population is expanded in the IPF lung and in an *ex vivo* PCLS assay, suggestive of progenitor cell behavior.

## 1. Introduction

Alveolar epithelial type 2 cells (AT2s) are surfactant-producing cells which serve as alveolar progenitors and play an essential role in the innate immune response. AT2s interact with macrophages through the secretion of various cytokines in response to pathogens and alveolar damage. As a result, AT2s activate macrophages and defend the alveolus [1][2]. Furthermore, AT2 interaction with Foxp3^pos^ Treg cells is critical for the repair of the epithelium during lung regeneration. Additionally, AT2s display enhanced proliferation when co-cultured with Foxp3^pos^ Treg cells [3]. However, our knowledge about the role of the interaction of AT2s with different immune cells in the repair of the epithelium or during pathogenesis of lung diseases remains limited [4][5][6].

AT2 heterogeneity has been demonstrated in different mouse models during homeostasis and regeneration/repair [7][8] [9,10]; however, the presence and function of the human equivalent of mouse AT2 progenitor cells are largely unknown [11][12]. Using *Sftpc^CreERT2/+^: tdTomato^flox/flox^* mice, and based on the differential level of the tomato reporter between two lineage-traced alveolar epithelial subpopulations, we have previously reported the existence of a novel population of AT2s, named Tom^low^ AT2s and which are enriched in programmed death-ligand1 (Pd-l1) [13]. This AT2 subpopulation expresses low levels of *Sftpc*, *Sftpb*, and *Sftpa1* and displays a low level of Fgf signaling activation. Following pneumonectomy, Tom^low^ AT2s are activated and display progenitor-like properties as they are amplified and exhibit elevated *Fgfr2b*, *Etv5*, *Sftpc*, *Ccnd1,* and *Ccnd2* expression compared to sham.

In addition, analysis of the behavior of these cells *ex vivo* in precision-cut lung slices (PCLS) from *Sftpc^CreERT2/+^: tdTomato^flox/flox^* mice supports their progenitor-like function in the context of significant injury to the AT2 lineage. Therefore, we named these Tom^low^ AT2s “injury-activated alveolar progenitors”, or IAAPs [14]. Pd-l1 is an immune inhibitory membrane receptor-ligand expressed in different immune and epithelial cells. Pd-l1 ligands bind to programmed death-1 (Pd-1) receptor. Pd-1 is expressed on a subset of Cd4^pos^ and Cd8^pos^ T cells, natural killer T cells, B cells, and activated monocytes. Pd-l1 is an immune inhibitory molecule that controls the inflammatory response to injury [15][16][17][18][19]. Remarkably, human PD-L1 expression is upregulated in non-small-cell lung cancer and adenocarcinoma. In the tumor microenvironment, cancer epithelial stem cells utilize the PD-L1 pathway to escape immune system surveillance by suppressing the cytotoxic response following binding of PD-L1 ligands expressed by cancer stem epithelial cells to the PD-1 receptor on T cells [20][21][22][23].

Interestingly, the presence of membrane PD-L1^pos^ alveolar and bronchiolar epithelial cells in a subgroup of IPF patients was previously reported. The presence of these cells is associated with higher fibroblast foci detection and patchy fibrosis compared to patients negative for PD-L1. However, no correlation between the expression of PD-L1 and honeycomb formation could be detected [24]. We also found in the normal human lung a subcluster of AT2 *PD-L1*^high^ cells following data mining of the scRNA-seq database [25]. These AT2 *PD-L1*^high^ cells express low levels of *ETV5*, *SFTPC*, *SFTPA1* and are enriched in the expression of immune system-related genes (*CXCL1*, *CXCL2*, *CXCL3*, *CXCL8*, *ICAM1* and *IRF1*) [13]. Intriguingly, an increased number of AT2 *PD-L1*^high^ cells expressing chemokines was also observed in the epithelium of patients with chronic obstructive pulmonary disease. These cells display higher expression of inflammation-related genes like *CXCL1*, *CXCL8*, *CCL2*, and *CCL20*, which supports a strong interaction of these cells with immune cells [26].

In this paper, we analyzed the available scRNA-seq dataset in the IPF cell atlas to investigate the presence of equivalent IAAP PD-L1^pos^ cells in human lungs. In particular, we quantified by flow cytometry the abundance of EpCAM^pos^HTII-280^neg^PD-L1^pos^ cells (potentially equivalent to the IAAPs), as well as EpCAM^pos^HTII-280^pos^PD-L1^neg^ cells (equivalent to mature AT2s) in IPF vs. donor human lungs. We also examined the expression of key genes in these populations by qPCR and monitored the growth of these cells in organoid and precision-cut lung slice assays. Our data indicate that the equivalent of IAAPs, initially identified in mice, exist in humans, and are differentially regulated in IPF vs. donor lungs.

## 2. Materials and Methods

### Human specimens

Human lung tissues from idiopathic pulmonary fibrosis (IPF) patients undergoing lung transplantation and non-IPF donors were collected in frame of the European IPF registry (euIPFreg) and provided by the UGMLC Giessen Biobank, member of the DZL platform biobanking. The study protocol was approved by the Ethics Committee of the Justus-Liebig-University School of Medicine (No. 111/08: European IPF Registry and 58/15: UGMLC Giessen Biobank), and informed consent was obtained in written form from each subject. Explanted lungs (n=9 for sporadic IPF) or non-utilized donor lungs or lobes fulfilling transplantation criteria (n=12; human donors) were obtained from the Dept. of Thoracic Surgery in Giessen, Germany and Vienna, Austria. All IPF diagnoses were made according to the American Thoracic Society (ATS)/European Respiratory Society (ERS) consensus criteria (American Thoracic Society Idiopathic Pulmonary Fibrosis: Diagnosis and Treatment International Consensus Statement, n.d. 343434), and a usual interstitial pneumonia (UIP) pattern was proven in all IPF patients.

### Mouse experiments

Animal studies were performed in accordance with the Helsinki convention for the use and care of animals and were approved by the local authorities at Regierungspräsidium Giessen (G7/2017-No. 844-GP and G11/2019-No. 931-GP).

### Human lung tissue dissociation

Subpleural lung tissue from explanted IPF lungs or excised from donor lungs due to donor-recipient size miss-match, was dissected and placed on ice. Tissue was minced with three scissors to pieces of approximately 1-2 mm in diameter, washed three times with RPMI and incubated for 30 mins at 37°C in RPMI containing Dispase (BD Cornig, 1:10 dilution) and 30 μg/μl DNase I with continuous and gentle agitation on a rotating shaker. Tissue pieces were first strained using surgical gauze, and then with 100 μm, 70 μm and 40μm cell strainers, and the strained solution was centrifuged and re-suspended in RPMI containing 10% FBS. Cells were washed two more times with RPMI 10% FBS, counted and shock-frozen in liquid nitrogen at a concentration of 2-10×10^6^ per vial until further use.

### Mouse lung tissue dissociation

Saline perfused lungs were removed from the thoracic cavity, and 1 ml RPMI solution containing 10% Dispase and 30μg/μl DNase I was instilled intra-tracheally and placed in a conical tube with 3ml of the same dissociation solution. Following 30 mins incubation in a 37°C water-bath, the lung lobes were isolated, minced and resuspended in 10 ml RPMI 10% FBS 30μg/μl DNase I. The lung suspension was then pipetted repeatedly up and down until the individual lung pieces consolidated in one single piece of floating extracellular material and the solution became turbulent. The resulting cell suspension was filtered through 70μm and 40μm cell strainers. Cells were then spun down two times at 1200 RPM for 5 min, resuspended in FACS-buffer (HBSS 2% FBS 10μg/ml DNase I, 2mM HEPES, 1% Penicillin Streptomycin), and placed on ice in preparation for flow cytometry staining.

### Flow cytometry and sorting

All analyses were performed on Li-nitrogen stored cell preparation available through the European IPF registry biobank. Human lung cell suspensions were thawed rapidly and cells were washed in 10 times the freezing volume of RPMI 10% FBS and maintained on ice throughout the procedure. Cells were resuspended in FACS buffer (HBSS, 2% FBS, 2μg/μl DNase I, 2mM HEPES, 1% Penicillin Streptomycin) at a concentration of 10^6^ cells/100μl and cell surface staining was performed using antibody master mixes whenever possible (antibodies in Table S1). For intracellular staining, cell surface staining was first performed, then cells were fixed and permeabilized for 30 mins at room temperature using the Foxp3/Transcription Factor Staining Buffer Set (eBiosciences), according to the manufacturer’s instructions. Cells were washed following fixation and resuspended in 100μl of permeabilization/wash buffer containing the primary antibody and incubated for one hour at room temperature, washed three times, followed by Anti-rabbit IgG (H+L), F(ab’)_2_ Fragment (Alexa Fluor^®^ 555 Conjugate Cell Signaling Technologies 1:1000).

The following controls were used: single color controls for instrument set-up, fluorescence-minus-one (FMO controls) for gating, and no primary controls for indirect intracellular staining.

Data were acquired on a BD FACSCanto II (BD Biosciences) using BD FACSDiva software (BD Biosciences). Data were further analyzed using FlowJoX software (FlowJo, LLC).

For the cell sorting experiments, samples were thawed successively and stained as above. Cells were sorted through a 100μm nozzle in RPMI 10% and the sort purity was assessed for each sample. The cells were immediately centrifuged and resuspended in lysis buffer (RNA Mini kit) for RNA isolation. The entire procedure was performed at 4°C and cell lysates were stored at −80°C until RNA isolation was performed.

### RNA extraction and quantitative real-time PCR

Following lysis of FACS-isolated cells from human and mouse lungs in RLT plus, RNA was extracted using a RNeasy Plus Micro kit (Qiagen) and cDNA synthesis was carried out using the QuantiTect reverse transcription kit (Qiagen) according to the instructions provided by supplier. Thereafter, selected primers were designed via NCBI’s Primer-BLAST option (https://www.ncbi.nlm.nih.gov/tools/primer-blast/, primer sequences in Table S2). Quantitative real-time polymerase chain reaction (qPCR) was performed using the PowerUp SYBR Green Master Mix kit according to the manufacturer’s protocol (Applied Biosystems) and a LightCycler 480 II machine (Roche Applied Science). *Hypoxantine‐guanine phosphoribosyltransferase* (*Hprt*) and *Glyceraldehyde 3-phosphate dehydrogenase* (*GAPDH*) were used as reference genes for mouse and human, respectively. Data were presented as mean expression relative to *Hprt* and *GAPDH*. Data were assembled using GraphPad Prism software (GraphPad Software, La Jolla/CA). Statistical analyses were performed utilizing two tailed-paired Student t-test. Results were regarded significant when p < 0.05.

### Microarray data

Differential gene expression was investigated using microarray analysis. Depending on the amount of RNA isolated per sample in an experiment, one of two possible microarray protocols was used. For RNA concentrations >50 ng/μl, the T7-protocol was followed. In this protocol, purified total RNA was amplified and Cy3-labeled using the LIRAK kit (Agilent Technologies, Waldbronn, Germany) following the kit instructions. Per reaction, 200 ng of total RNA was used. The Cy3-labeled DNA was hybridized overnight to 8×60K 60mer oligonucleotide spotted microarray slides (Agilent Technologies, design ID 028005). For experiments where samples yielded <50 ng/μl of RNA, the SPIA-protocol was utilized. In this protocol, purified total RNA was amplified using the Ovation PicoSL WTA System V2 kit (NuGEN Technologies, Leek, The Netherlands). Per sample, 2 μg amplified cDNA was Cy3-labeled using the SureTag DNA labeling kit (Agilent Technologies). The Cy3-labeled DNA was hybridized overnight to 8×60K 60mer oligonucleotide spotted microarray slides (Agilent Technologies, design ID 074809). For each protocol, hybridization, and subsequent washing and drying of the slides were performed following the Agilent hybridization protocol. The dried slides were scanned at 2 μm/pixel resolution using the InnoScan is900 (Innopsys). Image analysis was performed with Mapix 6.5.0 software, and calculated values for all spots were saved as GenePix results files. Stored data were evaluated using the R software (version 3.3.2) and the limma package (version 3.30.13) from BioConductor. Gene annotation was supplemented by NCBI gene IDs via biomaRt.

### Organoid assay

Sorted AT2 (CD45^neg^ CD31^neg^ EpCAM^pos^ HTII-280^pos^) and CD274^pos^ (CD45^neg^ CD31^neg^ EpCAM^pos^ HTII-280^neg^ CD274^pos^) cells from both donor and IPF lungs were centrifuged and resuspended in cell culture medium (RPMI Life Technologies, 10% FBS). Donor interstitial fibroblasts from one donor lung were provided by the Giessen Biobank [27] and were used as the mesenchymal component of the organoids. Mesenchymal (4000 cells/insert) and epithelial cells (5000 AT2s or 4000 CD274^pos^) suspensions were mixed in 100μl cell culture medium, followed by the addition of cold Matrigel^®^ growth factor-reduced (Corning) at a 1:1 dilution resulting in 200μl final volume per insert. Matrigel cell suspension was placed on the top of the filter membrane of the insert and incubated at 37°C for 30 min. Next, 350μl of the medium was transferred to each well. Cells were incubated under air-liquid conditions at 37°C with 5% CO2 for 12 days. Media were changed three times per week.

### PCLS culture and dissociation

One segment of each explanted human IPF lung was filled with 1.5% Low Melt Agarose (Bio-Rad) at 37°C and allowed to cool on ice for 30 mins. Blocks of tissue of ~3×3×4 cm (depth×width×height) were cut and prepared for sectioning using a Vibrating Blade Microtome (Thermo Fisher) at a thickness of 500μm. PCLSs were cultured for 3 days in DMEM/F12 (Gibco) supplemented with 10% human serum (Human Serum Off The Clot, Biochrom) and 1% penicillin/streptomycin in the presence of DAPT (10– 50μM in DMSO) or DMSO (1:1000; Sigma-Aldrich). The medium was changed daily. At the end of the experiment, PCLSs were dissociated using Dispase (1:10 dilution; Roche) and 10μg/μl DNase I for 45 mins at 37°C and processed for flow cytometry (see above).

## 3. Results

### 3.1 *CD274* mRNA expression in donor and IPF lungs

We previously identified a population of AT2s in mice and humans that express high levels of CD274 (also known as PD-L1) on their surface. In mice, these cells have quiescent progenitor characteristics. In humans, their role during homeostasis and in pathological conditions remains to be elucidated. To determine the mRNA expression level in the epithelial compartment at the single-cell level, two recently published scNGS data sets of donor and IPF lung cells (Kaminski & Banovich) were interrogated (Figure S1). In both data sets, *CD274* expression was found in a sub-population of AT2s as well as in IPF-specific aberrant basaloid and KRT5-/KRT17+ cells (Figure S1A). Although the expression pattern was concordant between the two data sets, they disagreed on the change in *CD274* expression level between the donor and IPF AT2s (Figure S1A). To determine the potential functional role of CD274 in pathogenesis, we searched the same databases for the expression of its cognate receptor *PD-1* (also known as the *PDCD1* gene) and found that it is specifically expressed in T-cells, and its expression was increased in IPF (Figure S1B). Thus, the data demonstrate that *CD274* is expressed in human alveolar epithelial cells and its expression is modulated in IPF. The complementary expression of its receptor on the surface of T-cells suggests CD274 may play a functional role in the alveolar epithelium-immune cell interaction. This result is supported by the previously reported high level expression of immune-related genes in IAAPs [13].

Next, focussing on the alveolar epithelial lineage (Figure 1A), we mined existing scRNAseq (Kaminski [28], Kropski [29] and Stripp [30], Figure 1B) for the presence of *CD274*^pos^ cells in donor lungs (Figure 1C), as well as in pathological lungs belonging to the interstital lung diseases (ILDs) spectrum, including IPF, NSIP, unclassifiable ILD, cHP and Sarcoidis (Figure 1D-H). Our results indicate that *CD274*^pos^ cells are detected in the alveolar lineage of donor and pathological lungs (Figure 1I). Comparison of *CD274*^pos^ cells in control vs. fibrotic cells (across all ILDs) indicates that *CD274* expression is significantly increased in fibrotic vs. control lungs (p=2.2 10^−16^) (Figure 1J). This difference was also significant for the other aforementionned diseases (Figure 1K) suggesting that *CD274* expression is increased in ILDs. We also reported in the mouse that *Fgfr2b*, *Etv5* and the differentiation markers *Sftpa*, *Sftpc* and *Sftpb* were upregulated in activated IAAPs following pneumonectomy [13]. This result suggested that in mice, there is a general activation of Fgfr2b signaling in IAAPs in the context of repair. This prompted us to investigate the status in human PD-L1^pos^ cells of the Fgfr2b signature identified in the mouse at embryonic day (E) 12.5 [31], E14.5 [32] and E16.5 [33]. In the adult lung, this Fgfr2b signature is mostly present in the alveolar epithelium and is found to be conserved between mice and humans ([33] and data not shown). Our results indicate that this signature is significantly increased in *CD274*^pos^ cells in IPF vs. control (Figure 1L-N), suggesting the reactivation of this AT2-specific developmental pathway in this population.

**Figure 1.**
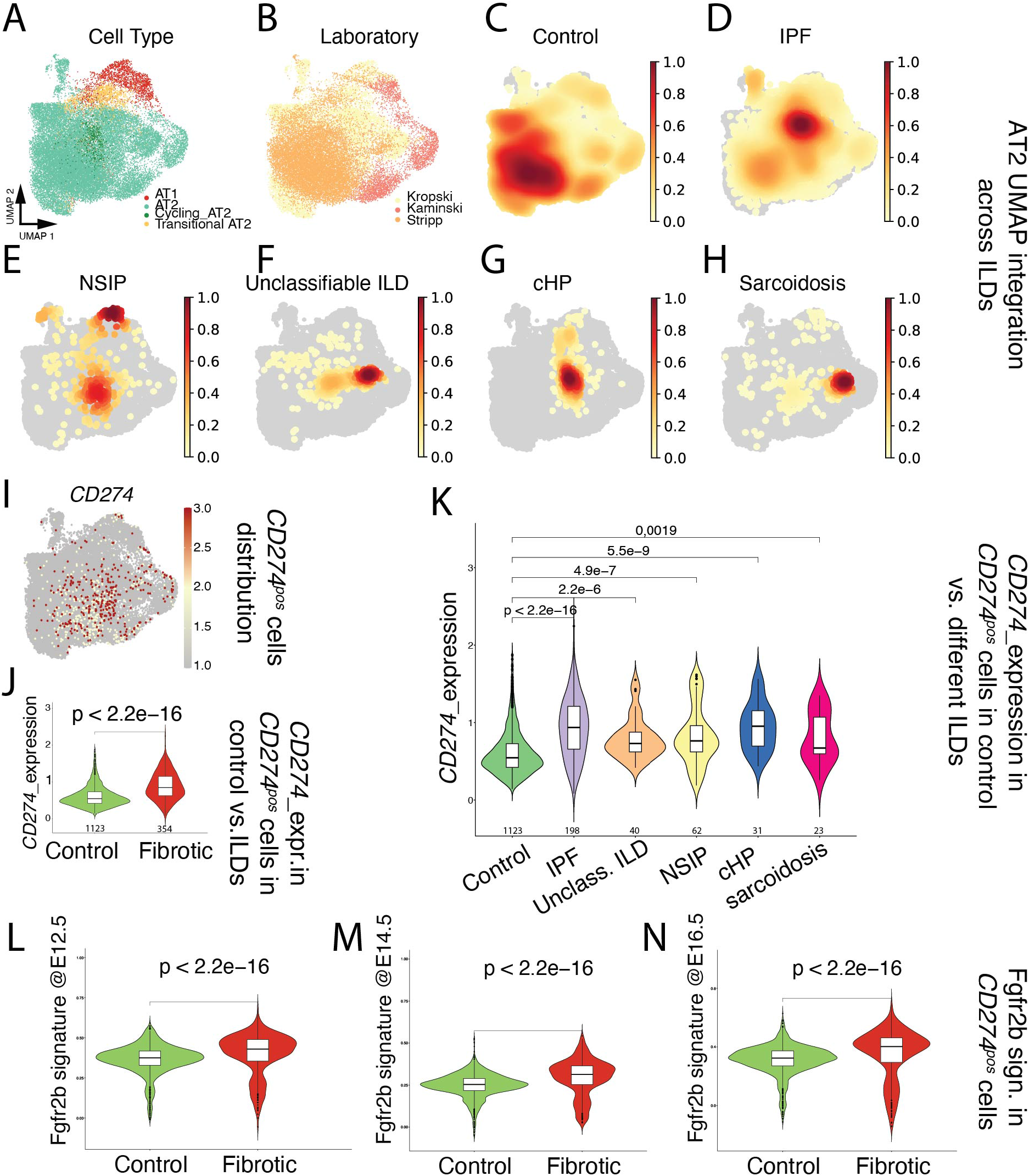
Single cell transcriptome analysis of distal epithelium in control and end-stage fibrotic donors. (A) Dimensional reduction of integrated data, showing cell type distribution, visualized by UMAP. (B) Dimensional reduction of integrated data derived from 3 published datasets, visualized by UMAP, with cells colored by lab of origin. (C-H) Distribution of cell density by disease. IPF: idiopathic pulmonary fibrosis, NSIP: non specific interstitial pneumonia, unclassifiable ILD: interstitial lung disease, cHP: chronic hypersensitivity pneumonitis. (I) Expression of *CD274* (*PD-L1*), visualized by UMAP. (J) Expression of *CD274* (*PD-L1*) in positive cells, comparing control and fibrotic datasets, visualized by violin plots. The value underneath each violin represents number of expressing cells. (K) Expression of *CD274* (*PD-L1*) in positive cells, comparing control and each fibrotic disease, visualized by violin plots. The value underneath each violin represents number of expressing cells. (L-N) Expression of mouse Fgfr2b signatures identified at E12.5, E14.5 and E16.5 on human *CD274* (*PD-L1*) positive cells, comparing control and fibrotic datasets, visualized by violin plots.

However, and as a note of caution, direct comparison of two key genes in the FGFR2b signature in humans in donor (n=4) and IPF (n=4) *CD274*^pos^ populations isolated for this study from our biobank revealed that IPF *CD274*^pos^ cells express lower levels of *FGFR2b* and *ETV5* compared to the corresponding expression in donor CD274^pos^ cells (Figure S2); this suggests important heterogeneity in terms of FGFR2b signaling activation in IPF vs. donor *CD274*^pos^ cells between different datasets. The significance of FGFR2b signaling in the *CD274*^pos^ cells is still unclear, and may be related to the capacity of these cells to display some of their progenitor-like characteristics, such as proliferation and/or differentiation.

### 3.2 Cell surface CD274 expression in donor and IPF lungs

To determine the functional pool of CD274, five donor and five end-stage IPF lung samples were dissociated into single-cell suspension and analyzed by flow cytometry. CD274 cell surface expression was analyzed in the epithelial compartment (CD45^neg^ CD31^neg^ EpCAM^pos^) as a function of HTII-280, an AT2 specific marker (Figure S3 A-C). Confirming previously published observations [34], the proportion of AT2s (Figure 2A, population Q1) was drastically diminished in IPF from an average of 80% in donors to approximately 25% in IPF epithelium. CD274 expression was not detected in AT2s, but a small HTII-280^neg^ CD274^pos^ population (called CD274^pos^ from here on, population Q3) was present in the donor and was statistically increased in the lungs of IPF patients (Figure 2A,B, Figure S3D, population Q3 donor vs IPF = 0.54% ± 0.255 vs 10.48% ± 10.32, respectively, p<0.001). Of note, the amount of cell surface CD274 was similar between donor and IPF cells, shown by the similar mean fluorescence intensity (MFI) of the Q3 population between the donor and IPF (Figure 2C, average Log(MFI) donor vs. IPF = 3.81 ± 0.08 vs. 3.92 ± 0.18, p=0.18).

**Figure 2.**
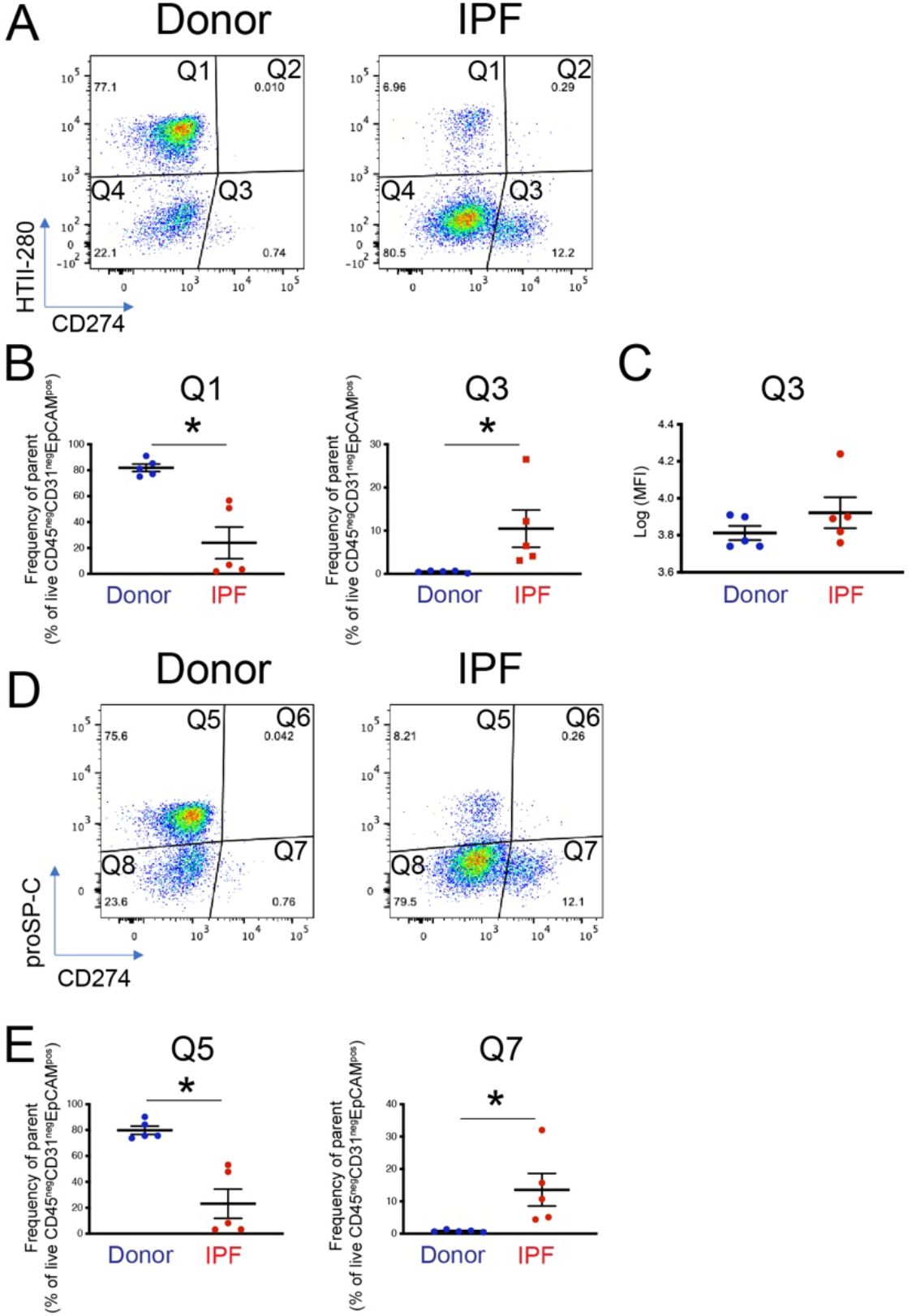
Cell surface expression of CD274 in donor and IPF human lung. (A) Representative flow cytometry panels of HTII-280 vs CD274 expression in the epithelial compartment (CD45^neg^ CD3^neg^ EpCAM^pos^) of donor and IPF lungs. (B) Quantification of the HTII-280 vs CD274 data shown in (A). (C) Quantification of the mean fluorescence intensity (MFI) of the CD274^pos^ (Q3 population) in donor and IPF samples. (D) Representative flow cytometry panels of proSP-C vs CD274 expression in the epithelial compartment (CD45^neg^ CD31^neg^ EpCAM^pos^) of donor and IPF lungs. (E) Quantification of the proSP-C vs CD274 data shown in (D). Statistical analysis was performed on log-transformed values, and student’s t-test was applied to determine statistical significance. * - p<0.05, ***-p<0.001.

In the mouse lung, CD274^pos^ cells express low levels of the AT2 marker proSP-C, and lineage tracing experiments show that they belong to the AT2 lineage. Thus, we analyzed the proSP-C and CD274 expression in the epithelial compartment of the same donor and IPF samples as in the previous analysis. Similar to the HTII-280 data, the number of proSP-C^pos^ cells was drastically decreased (Q5, donor vs IPF = 81.9% ± 7.05 vs. 23.9%± 25.1, p=0.014, Figure 2D,E and Figure S3E), and the number of CD274^pos^ cells was increased in the IPF lung (Q7 donor vs IPF = 0.84±0.34 vs 13.58±11.2, p<0.001). We also found that the CD274^pos^ population was entirely proSP-C negative (Figure 2D,E). Taken together, this data suggests that in the human lung, CD274 expression is confined to a population of HTII-280neg proSP-Cneg epithelial cells that are increased in the IPF lung.

### 3.3 Molecular characterization of the CD274^pos^ population in the mouse and human lungs

To gain insight into the identity of the corresponding CD274^pos^ population in the human lung, 4 donor and 4 IPF epithelial populations were sorted according to their HTII-280 vs CD274 expression pattern (Figure 3A, populations Q1 – HTII-280^pos^ CD274^neg^, Q3 - HTII-280^pos^ CD274^pos^ and Q4 - HTII-280neg CD274^neg^) and the expression of the *SFTPC* and *SCGB1A1* lineage-specific markers was determined in these isolated subpopulations by qPCR (Figure 3B). The Q1 and Q4 populations had a pattern of expression of *SFTPC* and *SCGB1A1* consistent with their respective alveolar and conducting airway lineages, while the Q3 population (CD274^pos^) cells expressed intermediate levels of *SFTPC*. The difference in the expression level of *SCGB1A1* between Q3 and Q1 subpopulations in either donor or IPF was not statistically significant (Figure 3B and C).

**Figure 3.**
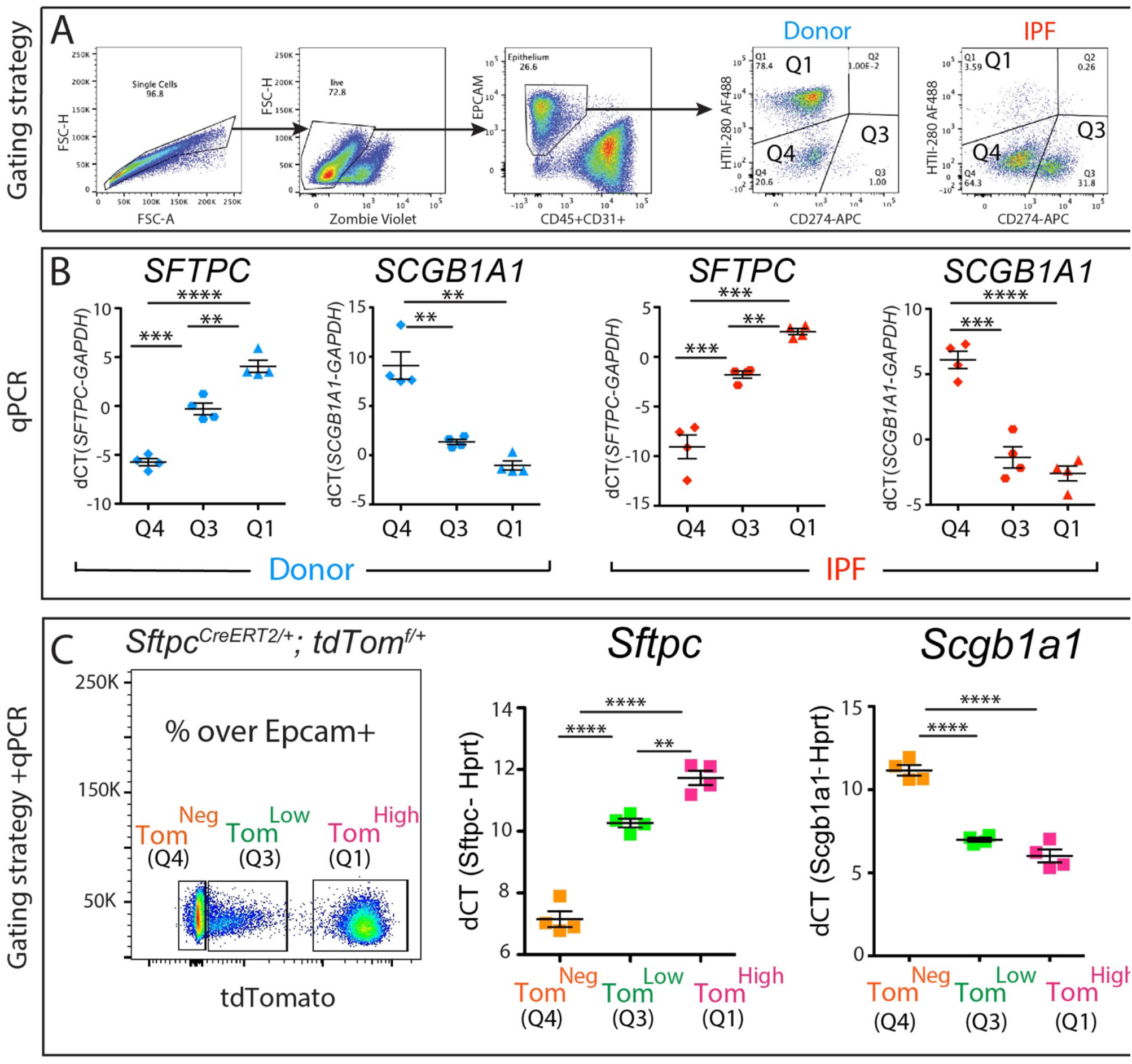
Molecular phenotype of the CD274^pos^ population. (A) representative sorting strategy to isolate HTII-280^pos^ CD274neg cells (Q1), HTII-280neg CD274^pos^ cells (Q3) and HTII-280^neg^ CD274^neg^ cells (Q4) in donors (n=4) and IPF (n=4). (B) Corresponding *SFTPC* and *SCGB1A1* mRNA quantification by qPCR in Q1, Q3 and Q4 subpopulations in donors (n=4) and IPF (n=4). (C) Representative flow cytometry panel of Tomato expression in the epithelial cell compartment of tamoxifen-treated *Sftpc^CreERT2/+^:tdTomato^flox/flox^* mice (left panel). qPCR analysis of *Sftpc* and *Scgb1a1* in the populations sorted according to the level of tdTomato expression (middle and right panel). Data are represented as means ± SEM. *p < 0.05, **p < 0.01, ***p < 0.001, ****p < 0.0001.

Our previously published data [13] led to the identification of a population of alveolar epithelial cells that express low levels of the lineage tracing fluorescence protein tdTomato (Tom^low^) alongside mature AT2s (Tom^high^ – Figure 3C). The Tom^low^ population is enriched in Cd274^pos^ cells and bears quiescent progenitor capabilities [13]. To understand the molecular phenotype of these populations, we performed qPCR analysis of *Sftpc* – an AT2 marker, and *Scgb1a1* – a club cell marker, on sorted Tomhigh (corresponding to Q1 subpopulation), Tom^low^ (corresponding to Q3 subpopulation) and Tomneg cells (corresponding to Q4 subpopulation) from tamoxifen-treated *Sftpc^CreERT2/+^: tdTomato^flox/flox^* mice. As expected, the Tom^high^ cells expressed high levels of the AT2 marker *Sftpc* and minimal levels of the club cell marker S*cgb1a1*, while the Tom^neg^ cells had the opposite expression pattern, conforming to their conducting airway identity. Interestingly, the Tom^low^ population had a level of *Sftpc* expression lower than mature AT2s but significantly higher than Tom^neg^, supporting our conclusion that Tom^low^ belong to the AT2 lineage (Figure 3C). Interestingly, Tom^low^ displayed a low level of expression of S*cgb1a1*, similar to what was observed in the Tom^high^ (Figure 3C).

Next, we carried out bulk RNA sequencing of isolated AT2s (Q1) and PD-L1 (Q3) subpopulations from n=2 donor and n=2 IPF patients. First, we explored the status of the AT2 signature in AT2s vs. PD-L1^pos^ cells in donor and IPF. Similar to what we found in the mouse [13], we found a significantly lower level of AT2 markers in PD-L1^pos^ cells vs. AT2s (Figure S4A). Interestingly, in our limited dataset, we did not observe any significant difference in the AT2 signature within PD-L1^pos^ cells in either donor or IPF. This result supports our previous observation that FGFR2b signaling, which is known to promote AT2 differentiation, is not increased in PD-L1^pos^ cells in IPF (Figure S2)

Next, we identified the most regulated pathways in AT2s and PD-L1^pos^ cells in IPF vs. donor (Figure S4B,C). In AT2s, the most dysregulated pathways were ribosome, focal adhesion, staphilococcus infection and chemokine signaling pathways. In PD-L1^pos^ cells, a very significant increase in the dynamic range of the changes observed between IPF and donor was noted compared to the the dynamic range of the changes observed in AT2s (−log_10_(p-value))= 23 vs. 12 in PD-L1^pos^ cells vs. AT2s, respectively). This result suggests tonic transcriptomic changes in the PD-L1^pos^ cells between donors and IPF. The most relevant dysregulated pathways in PD-L1^pos^ cells were pathways in cancer, cell cycle and Hippo signaling. This limited transcriptomic analysis, which should be expanded in the future, suggests that PD-L1^pos^ cells display a lower AT2 signature compared to the mature AT2s, and exhibit a dysregulation in the expression of genes linked to cell proliferation.

### 3.5 *In vitro* clonogenic potential

Mouse Tom^low^ cells combined with resident mesenchymal cells gave rise to fewer organoids in 3D culture compared to the AT2 population of Tom^high^ [13]. To study the *in vitro* clonogenic potential of the CD274^pos^ population, CD274^pos^ (Q3 population) and AT2s (Q1 population) were sorted from two donor patients and cultured in Matrigel in the presence of donor-derived interstitial fibroblasts and in a medium permissive of AT2 growth for 12 days (see Materials and Methods). In this assay, sorted AT2s gave rise to alveolospheres at the expected frequency (3-5%, Figure 4A). Although organoid formation was initiated in the CD274^pos^ population, the organoids failed to grow in the provided conditions (Figure 4A). However, counting the number of initiated colonies revealed a similar colony-initiating efficiency of this population, suggesting that these cells present colony initiation potential but failed to undergo proliferation (graph in Figure 4A).

**Figure 4.**
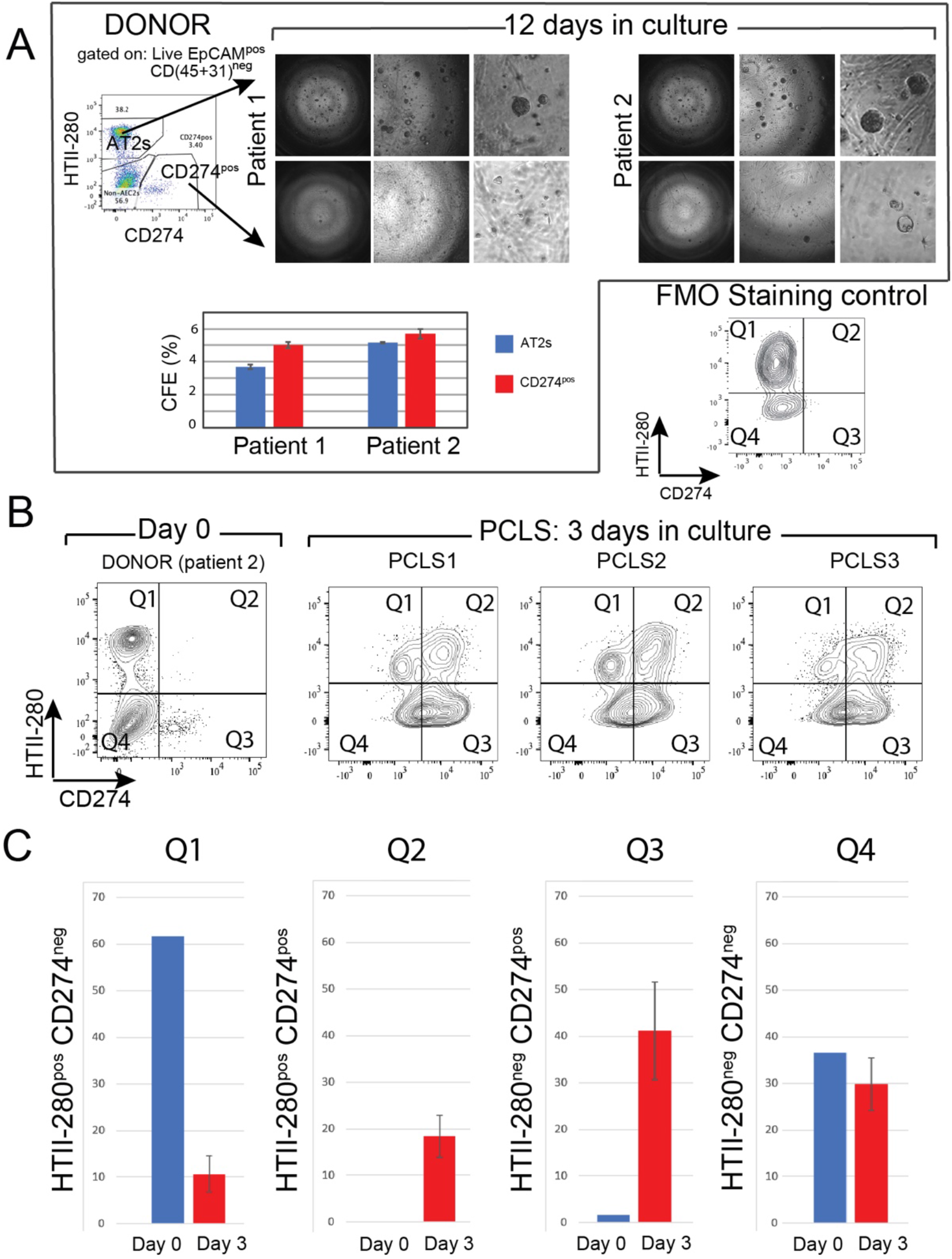
*In vitro* clonogenic potential of human CD274^pos^ epithelial cells. (A) Representative images of AT2- and CD274^pos^-derived organoids from two donor patients. Quantification of the CFE. (B) Flow cytometry panel of the day 0 analysis of HTII-280/CD274 expression in the epithelial compartment of patient 2 used for PCLS generation and culture. Flow cytometry analysis of HTII-280/CD274 expression in the epithelial compartment of PCLS (n=3) cultured for 3 days compared to the cells isolated at day 0 from the lung of patient 2. (C) Quantification of the data in (B).

### 3.5 *Ex vivo* progenitor potential

To study CD274 expression in a more physiological model, we generated precision-cut lung slices (PCLS) from one donor lung (patient 2, Figure 4A, B). After three days of culture, PCLSs were dissociated and CD274 expression was analyzed in the epithelial compartment of three cultured PCLSs (Figure 4B). Cells isolated from the same patient at the time of PCLS generation were used as Day 0 control (Figure 4A). Flow cytometry analysis showed an overall increase in CD274 expression in both HTII-280^neg^ (Q3) and HTII-280^pos^ (Q2) populations, suggesting the expansion of CD274^pos^ cells in *ex vivo* conditions (Figure 4C). In the absence of lineage tracing, it is difficult to determine progenitor-progeny relationships in this assay. However, the increase in proportion in both populations strongly suggests the proliferation and differentiation of the original CD274^pos^ population, which is a hallmark of progenitor cells.

Interestingly, using mouse PCLS to investigate the fate of lineage-traced alveolar epithelial cells, we also reported massive loss of mature AT2s, as well as amplification of the IAAPs [13]. The increase in the percentage of HTII-280^pos^CD274^pos^ cells after 3 days of culture could therefore represent the differentiation of CD274^pos^ cells to HTII-280^pos^ cells.

Taken together, our data demonstrate the existence of a CD274^pos^ population in the human lung, which is expanded in the IPF lung, and which has a similar phenotype and functional behavior with the recently identified IAAPs in the mouse lung.

## 4. Discussion

The interplay between the immune system and structural cell types in the normal and fibrotic lung is a subject of high research interest. Several studies have so far pointed at the role of the check-point molecule CD274 (PD-L1) in the mesenchymal compartment of IPF patients [35][36][37], while its role in the epithelial compartment remains to be established. We have previously shown that in the distal lung epithelium, CD274 expression is limited to an alveolar epithelial population with quiescent progenitor properties. In this study, we examined the cell surface expression of CD274 in the epithelial compartment of the human donor and IPF lung. Mining the published single cell NGS data (Kaminski/Rosas, Banovich, http://www.ipflcellatlas.org) of IPF lungs, we determined the *CD274* mRNA expression in AT2s, in aberrant basaloid, and in *KRT5-/KRT17+* epithelial cells. However, flow cytometry analysis of the functionally active, cell-surface expressed CD274 identifies a population with an intermediate AT2/conducting airway phenotype, and which is significantly increased in the lung of patients with IPF. This population bares similar molecular characteristics and *in vitro* behavior with the previously identified IAAPs [13]. Moreover, in an *ex vivo* model of donor PCLS culture, these cells are expanded, suggesting that they have self-renewal capacity.

Although CD274 expression has been acknowledged in the alveolar and bronchiolar compartments of the IPF lung [24], this is the first study that assesses its cell-surface expression, which is related to its functional immune-suppressive role. In the context of IPF and bleomycin-induced lung fibrosis, CD274 induction plays a role in the SMAD3/GSK3-mediated fibroblast to myofibroblast fate induction and collagen deposition [37][36], and in the IPF fibroblasts’ invasive properties [35][38]. In our study, although we detect low levels of CD274 on the surface of mesenchymal cells (CD45^neg^ CD31^neg^ EpCAM^neg^), we do not find any significant change in expression between donor and IPF single cell preparations (data not show). However, our cell isolation protocol uses mild enzymatic dissociation (dispase) which is optimized for the extraction and enrichment of viable AT2 cells. In these conditions, mesenchymal cells are under-represented, and harsher enzymatic dissociation protocols are necessary for their extraction, in particular from IPF tissue [39]. Thus, in our publication, we refrain from concluding any difference in CD274 expression in the mesenchymal compartment of IPF patients.

We and others have shown that in IPF, the expression of PD-L1’s cognate receptor PD-1 is increased on the surface of CD4+ T cells, strongly suggesting that the PD-1/PD-L1 (CD274) interaction plays an important functional role in the context of lung fibrosis (Supplemental Figure 1 and [36]). Indeed, PD-1 expressing CD4+ T cells secrete TGFβ and IL17A, which in turn activate fibroblast collagen 1 secretion via a STAT3-mediated mechanism [36]. In the mouse lung epithelium, expression of Cd274 is increased in a population marked by low expression of the Tomato AT2 lineage label (Tom^low^)[13], and this population actively replenishes the mature AT2 pool following alveolar injury mediated by *Fgfr2b* deletion (termed injury activated alveolar progenitors (IAAPs)) [14]. Similar to the IAAPs, human donor CD274^pos^ cells do not expand significantly in alveolar-permissive 3D cell culture conditions. However, a higher number of CD274^pos^ AT2 and non-AT2 cells emerged in cultured donor PCLS. Further experiments are needed to establish lineage relationships between various populations of CD274-expressing cells in this model, but our data clearly shows the expansion of this population.

Expression of *CD274* is noted at the mRNA level in AT2s in the previously published scNGS data (Kaminski & Banovich) (Figure 1), but consistent cell surface expression in this cell-type was seen only after PCLS culture, suggesting different levels of regulation at the mRNA and cell-surface protein level. Interestingly, *CD274* mRNA expression was also noted in a very small sub-population of AT2s (less than 0.45%), which expands in the lungs of chronic obstructive pulmonary dysplasia. This population has particular inflammatory properties, like pro-inflammatory cytokine secretion, but CD274 expression at the cell surface level in these cells has not been analyzed [26].

The functional role of CD274 expression in the lung epithelium remains an open question. It is well known that CD274 plays an important immuno-suppressive role in various cancers, conferring immune evading capabilities to the expressing transformed cells [16][40]. However, CD274 is expressed widely in healthy tissues, suggesting that it has an important functional homeostatic role. Indeed, a variety of tissues and cell types (testis, brain, eye) are preferentially protected from the immune system, a phenomenon known as immune privilege, a process in which CD274 plays an important role [41]. It also protects hematopoietic stem cells against premature removal from circulation [42], and marks a population of quiescent progenitors that regenerate Lgr5 expressing intestinal stem cells [43]. In the hair follicle, CD274 plays an important role in the progression of the hair cycle, potentially conferring protection to hair follicle stem cells located in the bulge area of the outer root sheath during the catagen phase [44]. Consistent with a role of CD274 in immune tolerance of healthy tissues, treatment with anti-PD-1/PD-L1 antibody for various malignancies results in serious side-effects like gastro-intestinal toxicity, skin rashes, alopecia [45], kidney, liver and lung injury [46].

Checkpoint inhibitors (CPI) are widely used in the treatment of non-small cell lung adenocarcinoma (NSCLC). However, pneumonitis is an important side-effect of CPI therapy (10-20% of patients) [47]. Moreover, the pre-existence of interstitial lung diseases before cancer diagnosis increases the risk of CPI-induced pneumonia, suggesting an important functional role of the population of PD-L1 expressing cells [19]. Our results point in the direction of, although they do not fully prove, an immune-protective role of a potential epithelial progenitor in the human lung, whose precise identity and lineage composition remains to be further elucidated. Several studies proposed that CPI therapy might be beneficial to limit the fibro-proliferative reaction in IPF [35][38], but our data, together with the clinical studies mentioned above, suggest that such interventions, unless specifically targeted, might further injure the already precarious lung epithelial compartment in IPF.

## Supplementary Materials

The following are available online at www.mdpi.com/xxx/s1

## Author Contributions

## Institutional Review Board Statement

The study was conducted according to the guidelines of the Declaration of Helsinki and was approved by the Ethics Committee of the Justus-Liebig-University School of Medicine (No. 31/93, 29/01, and No. 111/08: European IPF Registry), and informed consent was obtained in written form from each subject. Animal studies were performed in accordance with the Helsinki convention for the use and care of animals and were approved by the local authorities at Regierungspräsidium Giessen (G7/2017-No.844-GP and G11/2019-No. 931-GP)

## Informed Consent Statement

Informed consent was obtained from all subjects involved in the study prior to biospecimen collection.

## Data Availability Statement

Transcriptomic data is publicly available at the following location: Microarray data for the isolated AT2s and PD-L1^pos^ cells from donor and IPF lungs are available under the accession number GSE195770.

## Acknowledgments

The authors would like to thank Gabriela Dahlem (Giessen Biobank) and Gabriela Michel (FACS core facility)

## Supplementary Tables

**Table S1:** Antibodies used for flow cytometry staining

| Primary antibody target | Company | Cat no | FACS (per 10*6 cells) |
| --- | --- | --- | --- |
| anti human proSP-B | Millipore | AB3430 | 1_200 |
| anti human EpCAM APC-Cy7 | Biolegend | 324222 | 0.5_100 |
| anti human CD45 Pe-Cy7 | Biolegend | 304016 | 2.5_100 |
| anti human CD31 Pe-Cy7 | Biolegend | 303120 | 0.5_100 |
| anti human HTII-280 | TerraceBiotech | TB-27AHT2-280 | 1_500 |
| anti human CD274 | Biolegend | 374514 | 1_100 |
| Donkey anti-rabbit Alexa-Fluor 488 | ThermoFisherScientific | A21206 | 1_500 |
| anti mouse EpCAM Pe-Cy7 | Biolegend | 118216 | 0.25_100 |
| anti mouse CD45 APC-Cy7 | Biolegend | 103114 | 0.25_100 |
| anti mouse CD31 APC-Cy7 | Biolegend | 102418 | 0.25_100 |
| anti mouse CD274 unconjugated | Thermo-Fischer | PA5-20343 | 1_100 |
| Goat anti rabbit antibody Alexa flour 488 (Invitrogen, 1:500) | ThermoFisherScientific | A11008 | 1_500 |

**Table S2:**
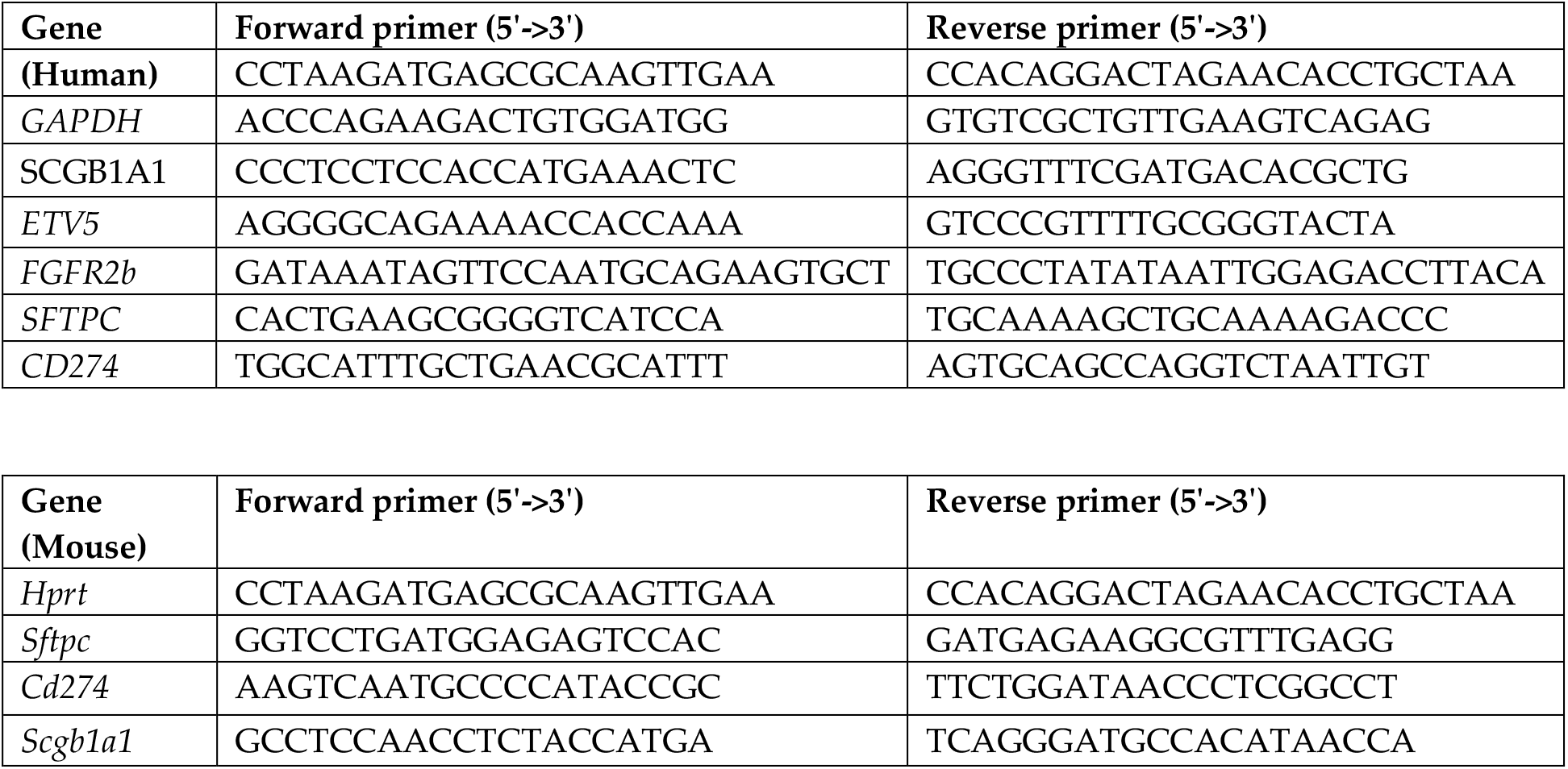
qPCR primers

## Supplementary figures

**Figure S1.**
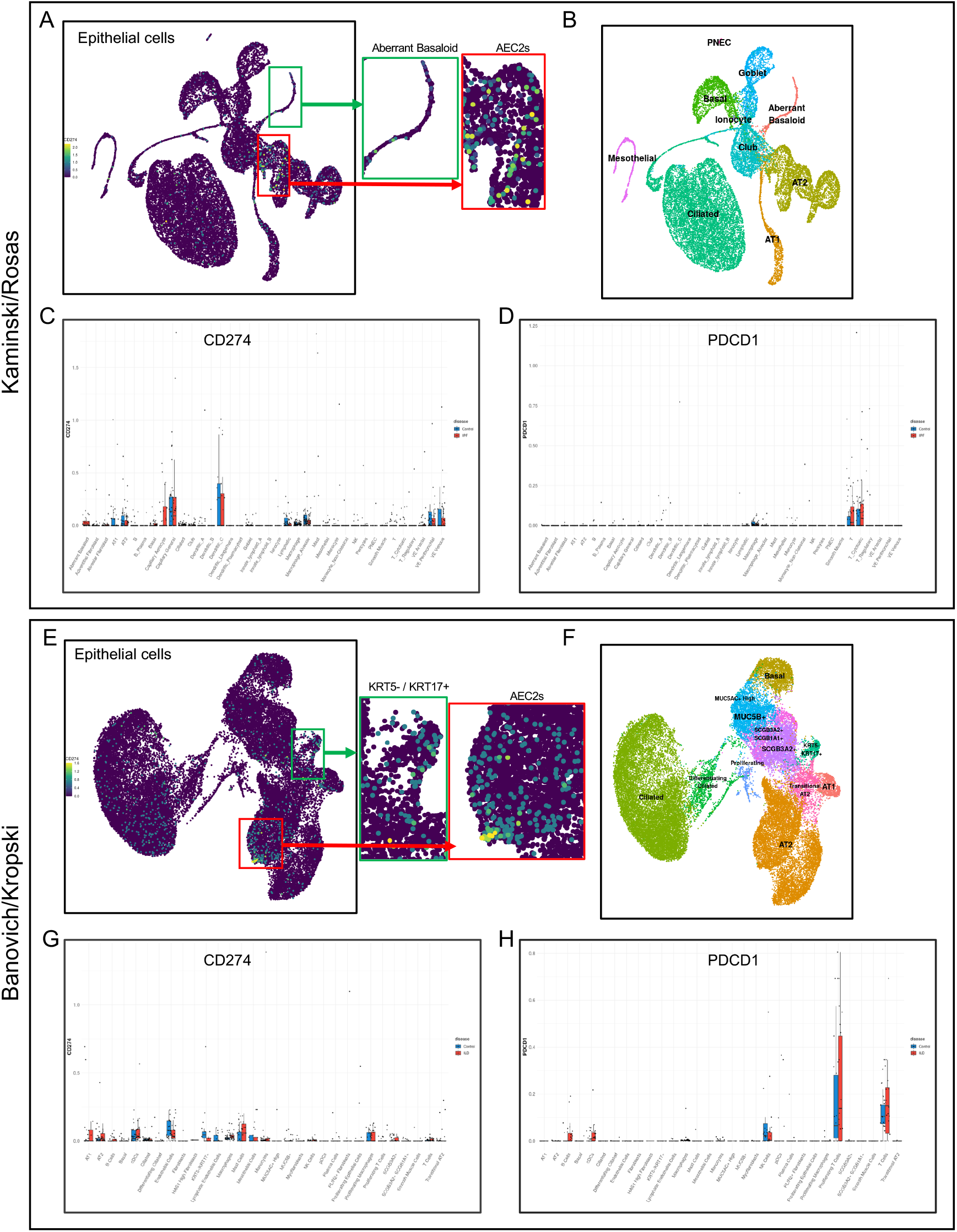
Expression of CD274 mRNA in donor and IPF lungs. Alveolar epithelium expression of *CD274* mRNA in the Kaminski/Rosas (A) and Banovich/Kropski (E) data sets shown as UMAPs. (B, F) UMAP representation of different epithelial cell type clusters in the Kaminski/Rosas (B), and Banovich/Kropski (F) data sets. (C, G) Overview of the *CD274* mRNA expression levels in donor and IPF throughout all lung cell types. (D, H) Expression of *PDCD1* (*PD1*) in the immune cell compartment of donor and IPF lungs in the Kaminski/Rosas (D), and Banovich/Kropski (H) data sets. Data retrieved from http://www.ipfcellatlas.com.

**Figure S2.**
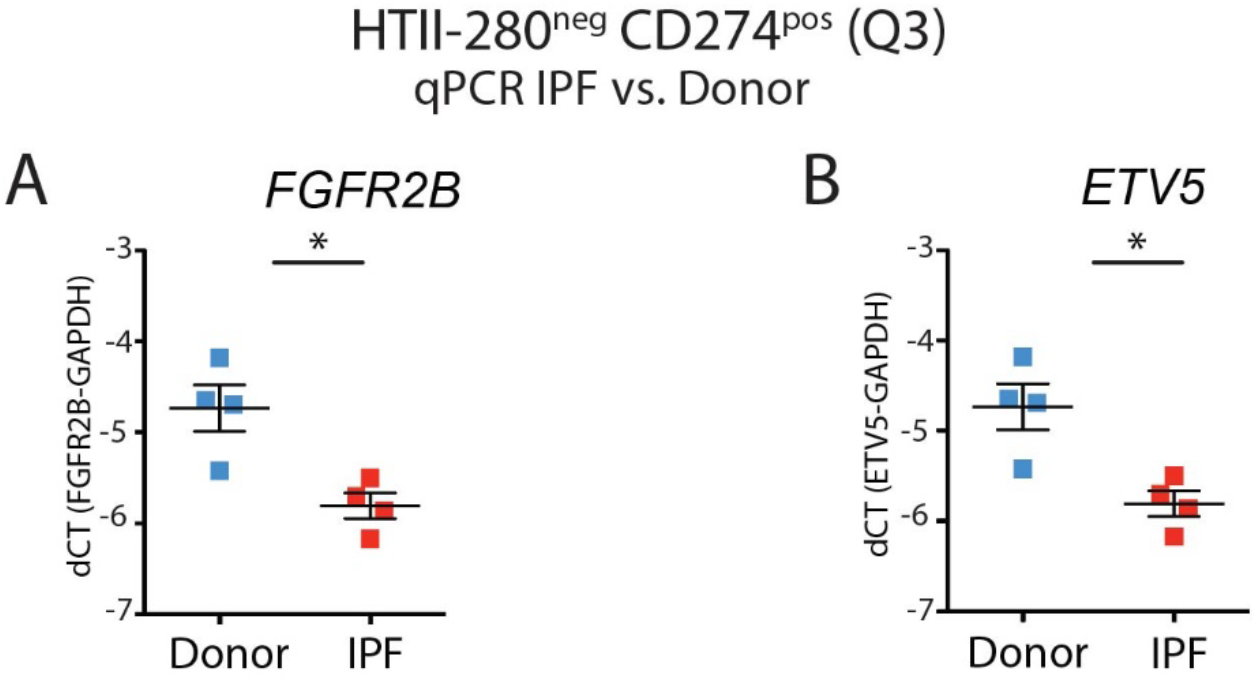
Expression of FGFR2B and ETV5 in donor and IPF-derived CD274^pos^ cells. (A) qPCR analysis of *FGFR2B* in HTII-280^neg^ CD274^pos^ cells (Q3) subpopulations in donors (n=4) and IPF (n=4). (B) qPCR analysis of *ETV5* in HTII-280^neg^ CD274^pos^ cells (Q3) subpopulations in donors (n=4) and IPF (n=4). Data are represented as means ± SEM. *p < 0.05, **p < 0.01, ***p < 0.001, ****p < 0.0001.

**Figure S3.**
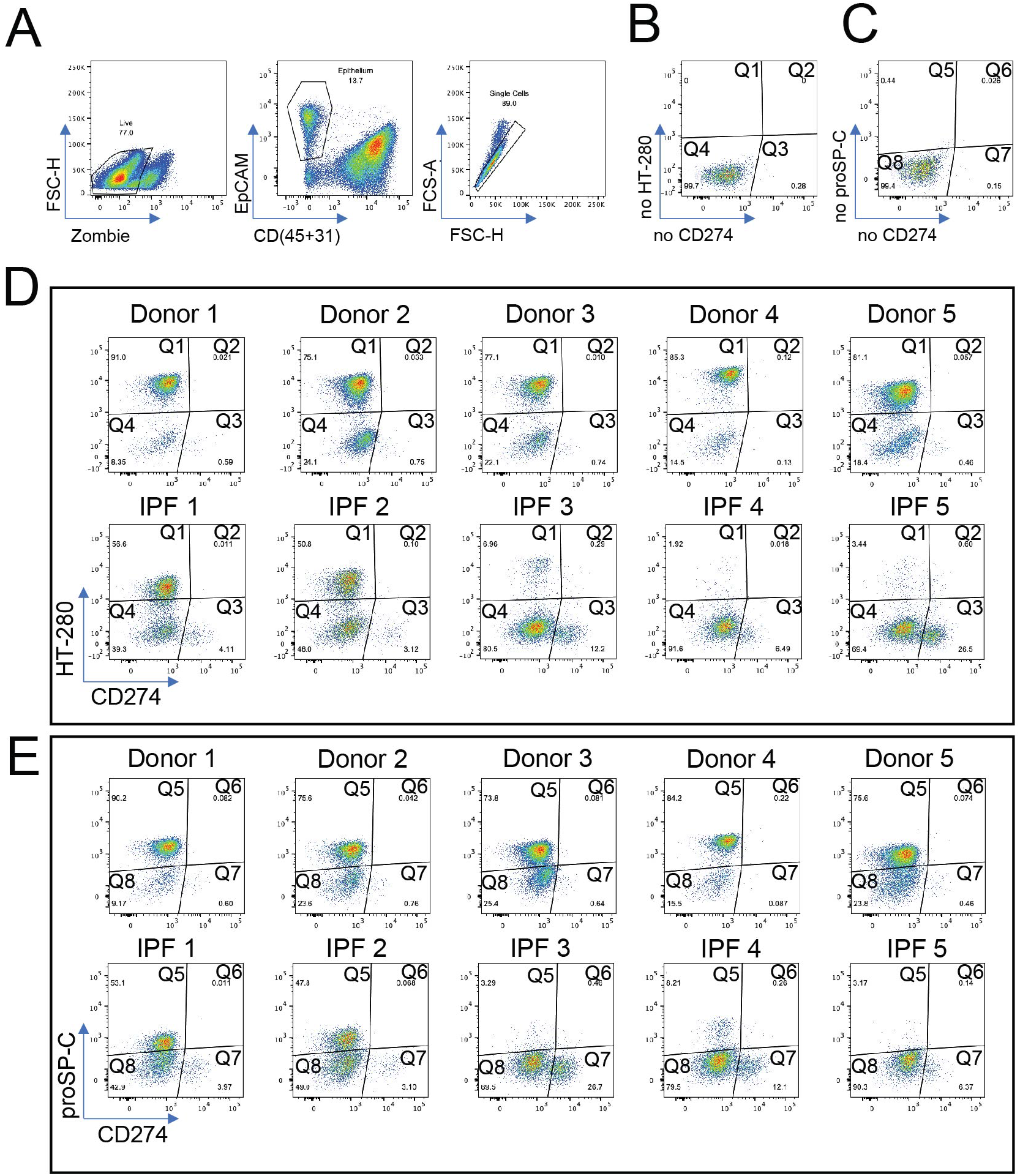
Cell surface expression of CD274 in donor and IPF human lung. Five donor and five end-stage IPF lung samples were dissociated into single cell suspension and analyzed by flow-cytometry. (A) Representative flow cytometry panels showing the gating strategy of the epithelial cells analyzed in Figure 2. First, live cells were identified as Zombie^neg^ cells (left panel), then epithelial cells were identified as CD45^neg^ CD31^neg^ EpCAM^pos^ (middle panel) and doublets were excluded based on FSC-A vs FSC-H analysis (right panel). (B) Controls where HTII-280 and CD274 (left panel) and proSP-C and HTII-280 were omitted were used to define the gating for the flow cytometry analysis in Figure 2. (D) Individual panels for each of the patients in the HTII-280 CD274 analysis. (E) Individual panels for each of the patients in the proSP-C CD274 analysis.

**Figure S4.**
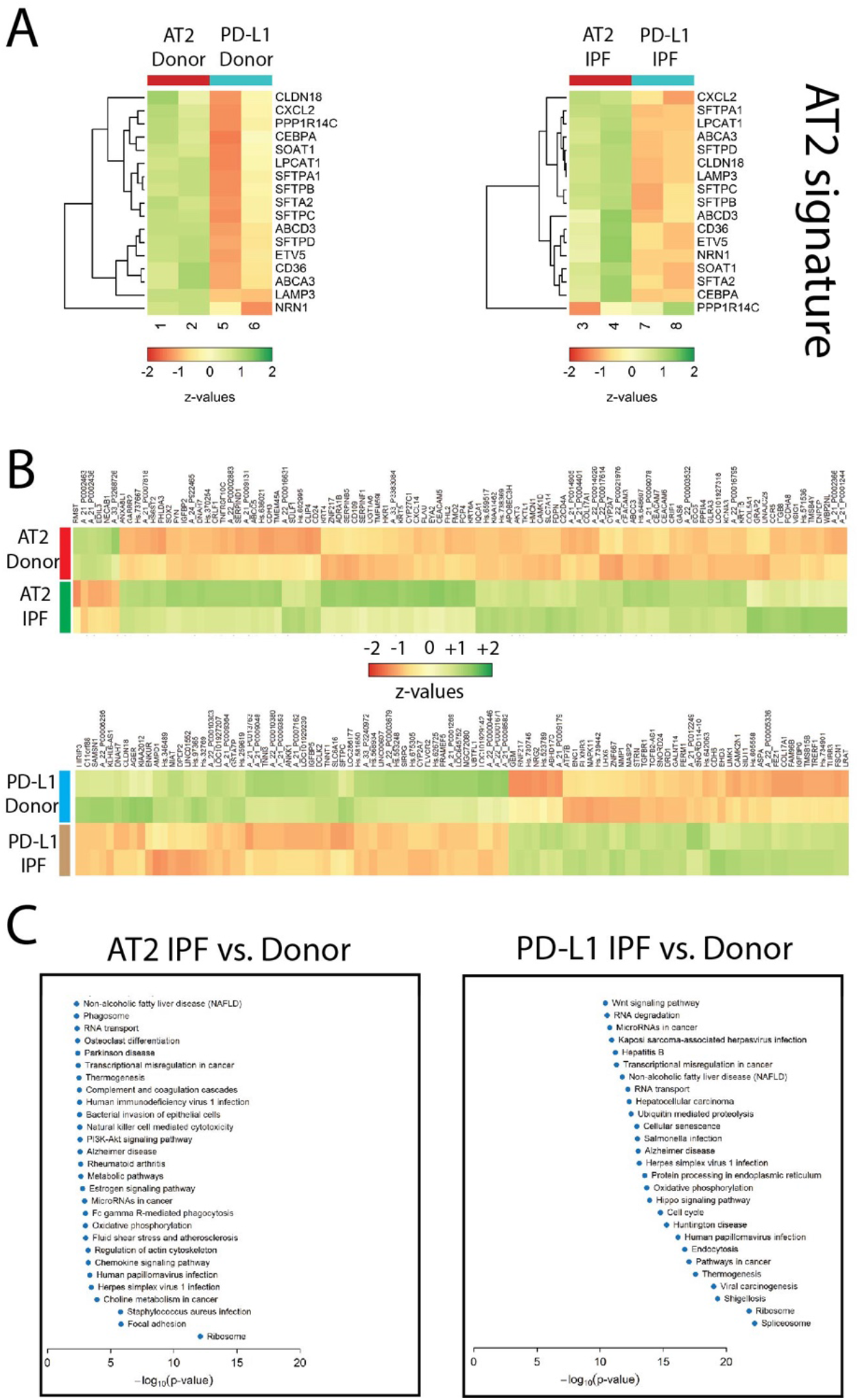
Transcriptomic analysis of donor and IPF-derived AT2 and CD274^pos^ (PD-L1^pos^) cell populations. (A) Heatmaps of AT2 cell signature genes based on the microarray data of isolated AT2 and PD-L1^pos^ cells in both donor (n=2) and IPF (n=2) lungs. (B) Heatmap of the top 100 differentially regulated genes in AT2 cells (according to the p-value) in donor vs. IPF, and heatmap of the top 100 differentially regulated genes in PD-L1^pos^ cells (according to the p-value) in donor vs. IPF. (C) Corresponding KEGG pathway analysis showing the top regulated pathways in AT2 and PD-L1^pos^ cells in IPF vs. donor, according to significance (−log^10^(p-value)).

